# Beyond distance-decay: The Mississippi River shapes gut microbiome communities in the *Peromyscus maniculatus* species complex

**DOI:** 10.64898/2026.06.03.729934

**Authors:** Danielle M. Blumstein, Aishwarya Patel, Gabriele Schiro, Taichi A. Suzuki

## Abstract

Biogeographic barriers are fundamental in shaping the distribution of animals and plants, yet the role of discrete landscape features in constraining microbial dispersal remains poorly understood. Identifying the barriers that partition gut microbial communities is essential for modeling the distribution of hosts and their symbionts across physical space and evolutionary time. The deer mouse (*Peromyscus maniculatus* species complex), which inhabits nearly all terrestrial environments in North America, provides an ideal model for testing these biogeographic drivers of the mammalian gut microbiome. Using NSF NEON biorepository samples, we characterized the gut microbiomes of 13 populations across the contiguous United States using full-length 16S rRNA long-read sequencing. While we observed a consistent distance-decay relationship across the continent, our results reveal that the Mississippi River acts as a major biogeographic break, significantly increasing microbial dissimilarity beyond the levels predicted by geographic distance alone. This “river effect” suggests that large fluvial systems impose a discrete barrier to microbial transmission, likely due to restricted host dispersal. Furthermore, we identified specific microbial lineages that exhibit differential sensitivity to this barrier, suggesting a gradient in microbial acquisition patterns ranging from the environment to host-host transmission. Together, these findings demonstrate that the gut microbiome may act as a sensitive bio-indicator of landscape-level ecological connectivity, revealing that large-scale landscape barriers disrupt microbial transmission even among closely related host populations that lack reproductive barriers.

## Introduction

Biogeographic barriers such as mountain ranges, rivers, and oceans play a fundamental role in shaping the distribution and diversification of plants and animals (Emerson et al., 2011; Ricklefs & Jenkins, 2011). The impact of isolation depends heavily on the ecological traits and dispersal capacities of the organism (Mazel et al., 2017; Peixoto et al., 2018; Procheş, 2006; Williams et al., 2024). However, the role of discrete landscape features in constraining microbial dispersal remains poorly understood. Understanding how geography structures host-associated microbial communities is important for several reasons. First, microbial dispersal influences the assembly and stability of host-associated microbiomes (Custer et al., 2022; Marrec & Lehtinen, 2026; Moeller et al., 2017; Nemergut et al., 2013), which in turn affect host physiology, immunity, and disease susceptibility (Chu et al., 2025; McFall-Ngai et al., 2013; Nouri et al., 2022; Suzuki, 2017). Second, identifying barriers to microbial movement has implications for disease ecology, including the spread of pathogenic or beneficial microbes among host populations (Chiu et al., 2017). Finally, understanding microbial taxa and traits associated with dispersal capability provides insight into the mechanism underlying microbial transmission in natural populations (Martiny et al., 2006; Nemergut et al., 2013).

A growing body of work shows that geography can explain variation in gut microbiomes across natural populations of mammals, including mice (e.g. Goertz et al., 2019; Linnenbrink et al., 2013; Suzuki et al., 2020), primates (e.g. Grieneisen et al., 2019; Moeller et al., 2013), and humans (e.g. Suzuki et al., 2022; Truong et al., 2017). These studies commonly report continuous distance–decay or isolation-by-distance patterns in overall microbiome similarity, where microbiomes become more dissimilar as geographic distance increases (Dillard et al., 2026; Tuoliu et al., 2025). In addition, certain gut microbial taxa show geographically restricted distributions, such as human population–specific strains observed across continents (Andreu-Sánchez et al., 2025; Suzuki et al., 2022). However, the generality of these community-level and microbial strain-level patterns across mammalian systems remains unclear.

Several challenges limit our ability to test how geographic barriers influence microbiome dispersal. Most microbiome studies rely on short-read sequencing of the 16S rRNA hypervariable regions, which often lack the resolution needed to detect strain-level transmission patterns (Buetas et al., 2024; O’Sullivan et al., 2025). Additionally, studies in non-human systems frequently compare different host species across deep evolutionary timescales (Chu et al., 2025; Groussin et al., 2017; Kohl et al., 2022; Li et al., 2025; Tuoliu et al., 2025), making it difficult to separate the effects of host divergence from geographic barriers. Finally, most host species have limited geographic ranges (Jensen et al., 2025; IUCN 2025), restricting the ability to test how landscape features shape microbiome structure across broad spatial scales. As a result, whether discrete landscape features create stepwise changes in microbiome composition, and which microbial taxa are most sensitive to such barriers, remains largely untested.

The deer mouse (*Peromyscus maniculatus* species complex) provides an ideal system to address these questions. Deer mice are among the most abundant and widely distributed mammals in North America, occupying a broad range of habitats across the continent (Boria et al., 2026; Bradley & Lindsey, 2019; Dragoo et al., 2006; Gozashti et al., 2026; Greenbaum et al., 2019; Kalkvik et al., 2012; Natarajan et al., 2015). Recent studies have revealed substantial population structure within this complex (Boria et al., 2026; Bradley & Lindsey, 2019; Finkbeiner et al., 2024; Natarajan et al., 2015). This group includes at least six geographically structured clades based on mitochondrial DNA (Dragoo et al., 2006; Kalkvik et al., 2012; Natarajan et al., 2015) and six highly supported lineages based on phylogenomic and population genomic analyses of whole genomes (Boria et al., 2026). Notably, the Mississippi River has been identified as a major barrier to gene flow separating eastern and western populations (Boria et al., 2026; Finkbeiner et al., 2024). Much of this structure appears to have formed during a rapid range expansion following the end of the Last Glacial Maximum (Boria et al., 2026). Despite this genetic structure, individuals from different clades can produce fertile offspring in captivity (Bedford & Hoekstra, 2015, Peromyscus Genetic Stock Center Columbia, SC, USA), indicating relatively recent divergence. This combination of broad geographic distribution, known host genetic structure, and relatively shallow evolutionary divergence makes deer mice a powerful system for testing how landscape barriers influence gut microbial dispersal across continental scales.

Here, we leverage samples collected by the National Ecological Observatory Network (NEON) from across the contiguous United States and full length 16S rRNA gene sequencing using long-read technology to test how geographic barriers shape gut microbiome structure in deer mice. The NEON sampling framework provides standardized collections across broad spatial scales, allowing us to compare populations on either side of major landscape features. In addition, full length16S rRNA gene sequencing improves taxonomic resolution, enabling more precise detection of microbial lineages (Buetas et al., 2024) that may exhibit geographically structured distributions. Using this approach, we test whether discrete landscape barriers produce stepwise shifts in gut microbiome composition and identify microbial taxa that show differential sensitivity to geographic barriers.

## Methods

### Sample Collection, DNA Extraction, and Sequencing

Samples collected by the National Ecological Observatory Network (NEON) as part of the NEON small mammal box trapping protocol were selected from the NEON Biorepository at Arizona State University (ASU) based on several requirements (Supplemental Table 1). Samples were assigned to a region, east or west, based on their geographic location relative to the main channel of the Mississippi River. To maximize genetic, geographical, and ecological variation, locations were separated at least 200km from each other. These included locations in California, Colorado, Kansas, New Hampshire, North Dakoda, Oklahoma, Utah, Tennessee, Virgina, Washinton, and Wisconsin (Supplemental Table 2). Individual collection efforts that included both host DNA and fecal vouchers as well as weight data were selected. Host genomic DNA was extracted from the tissue on a Biomek FX liquid handling station using a semi-automated DNA extraction protocol (Ivanova et al., 2006). Cytochrome Oxidase Subunit 1 5’ region (Reverse H9375 (5’-3’): ACTAAGAGAGTAGGATCCTCATCAATA and Forward P. mani-F-9197 (5’-3’): GGAATTTATGGGTCTACATTC) was used for *P. maniculatus* species identification at the Centre for Biodiversity Genomics.

Fecal samples were briefly thawed at room temperature before DNA extraction with total genomic DNA extracted using Zymo Quick-DNA Fecal/Soil Microbe Miniprep Kit. Extracts were sent to Desert Southwest Genomics Center at AUS in Tucson, AZ for Kinnex 16S library preparation and full-length 16S rRNA gene sequencing using the PacBio platform (Revio SMRT Cell) using the universal bacterial 16S primers 27F (forward: AGRGTTYGATYMTGGCTCAG) and 1492R (reverse: AAGTCGTAACAAGGTARCY).

### Computation Analysis and ASV Inference

All the code used to analyze the data is located at the GitHub repository (https://github.com/DaniBlumstein/pema_NEON). Raw sequencing reads were processed using the pb-16S-nf Nextflow pipeline (v0.6, https://github.com/PacificBiosciences/HiFi-16S-workflow) with default parameters to reorients the sequences, removes forward and reverse primers, filter, trim the sequences by length and average quality, and infer ASVs and assign taxonomy. QIIME2 (Estaki et al., 2020) (2024.2) and DADA2 (Callahan et al., 2016) were used with default parameters to denoise into amplicon sequence variant (ASVs).

### Statistical Analysis

The ASV table rarefied to 3,771 reads/sample, taxonomy assignments, and rooted tree outputs from QIIME2 (Estaki et al., 2020) were imported into R (R Core Team, 2025) (4.5.2) using both phyloseq (McMurdie & Holmes, 2013) (1.52.0) and MicrobiotaProcess (Xu et al., 2023) (1.23.0), allowing for all downstream statistical analyses to be performed in R (R Core Team, 2025) and visualizations using ggplot2 (Wickham, 2016). Beta-diversity of the microbiome was calculated using Bray-Curtis dissimilarity (BCD) and Sørensen dissimilarity, and principal coordinates analysis plots were generated using MicrobiotaProcess (Xu et al., 2023). A PERMANOVA was used to quantify multivariate community-level differences using the adonis2 function in the vegan package. Separate models were constructed for each taxonomic level (strain, species, and genus), and each model included region, latitude, and longitude as predictors.

Pairwise comparisons were then categorized into three groups: within west, within east, and between region (east vs west), allowing us to explicitly test whether microbiome similarity differs across the river relative to within-region expectations. Mantel tests were used to test for correlation between BCD or Sørensen dissimilarity and geographic sampling distance (km), calculated with Geosphere using the Haversine formula, with the package cultevo (1.0.2) for sample pairs within the eastern region, within the western region, and between region (east vs west) for BCD or Sørensen dissimilarity and geographic sampling distance (km) for samples pairs.

Lastly, we identified the top ten most abundant genera from our phyloseq object and summarized how often each ASV occurred in eastern versus western region samples. From these counts, we calculated a regional bias score for each ASV based on its relative proportion in each region, where −1 indicates the ASV is found exclusively in the west, +1 indicates it is found exclusively in the east, and 0 indicates the ASV is present in both regions. This score reflects the direction and strength of regional association for each ASV. We then tested the phylogenetic trees for the top ten genera for phylogenetic clustering using our regional bias score with the standard measure, Pagel’s λ (Lin & Peddada, 2020) using the phylosig function in phytools (Blomberg et al., 2003). Limiting the analysis to the top abundant genera was necessary to ensure sufficient power to quantify phylogenetic clustering of ASVs on bacterial phylogenies. We also performed an analysis of compositions of microbiomes with bias correction (ANCOM-BC) (Revell, 2024) to identify differentially abundant ASVs associated with the eastern region and the western region, which helped us identify candidate ASVs to examine in phylogenetic trees.

## Results

### Gut microbiome composition differs between eastern and western regions of the Mississippi River

Full-length 16S rRNA sequencing using PacBio Revio resulted in a median of 7,4973.5 post-QC reads per sample, and the total number of ASVs after rarefaction was 26,365. Microbial communities were dominated by *Bacillota* (72.8%), *Bacteroidota* (22.4%), *Desulfobacterota* (1.72%), and *Campilobacterota* (1.06%).

Principal coordinate plots of Bray-Curtis dissimilarity (BCD) (Figure 1B) and Sørensen dissimilarity (Figure 1C, Supplemental Figure 1) showed striking microbiome separation between eastern and western regions, with Mississippi River as a geographic boundary. The PERMANOVA models included region, latitude, and longitude as predictors and were significant at both metrics at the ASV, species, and genus levels (Supplemental Table 3). Although the sample size is small, populations from California within the western region showed distinct clustering from the other populations (Figure 1A, Supplemental Figure 1).

**Figure 1.**
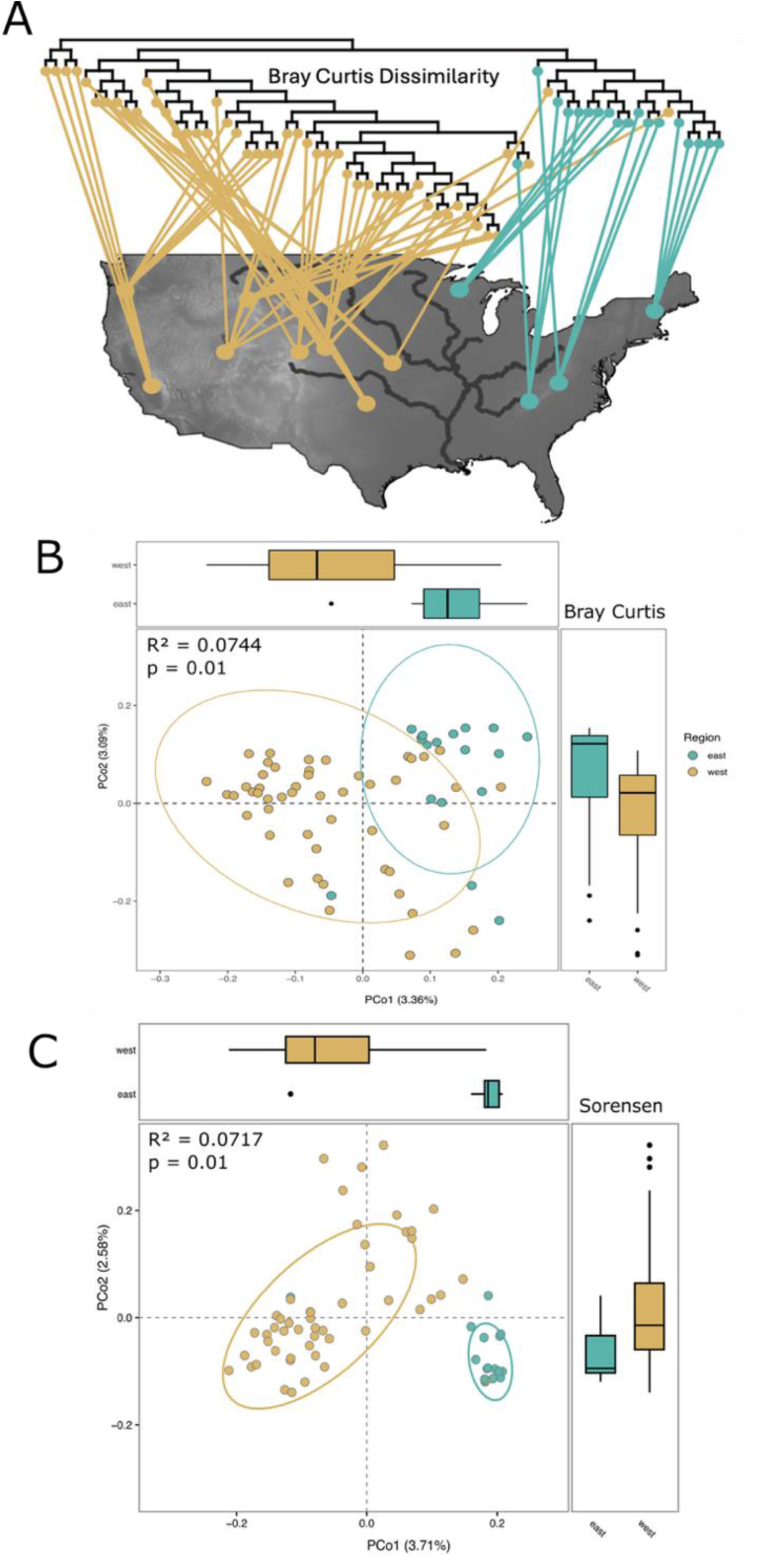
The relationship between sampling region and the gut microbiome of *Peromyscus maniculatus* (A) Bray-Curtis dissimilarity matrix projected onto the sampling locations of *Peromyscus maniculatus* across the North American distribution. (B) PCoA plot showing the effect of region on Bray-Curtis dissimilarity (PERMANOVA: R² = 0.0744, F = 1.6618, p = 0.01). (C) PCoA plot showing the effect of region on Sørensen dissimilarity (PERMANOVA: R² = 0.0717, F = 1.5957, p = 0.01). Colors correspond to samples collected on the western side of the Mississippi River (yellow) and the eastern side of the Mississippi River (blue).

The number of ASVs per sample ranged from 130 – 1132, and median ASV richness was significantly higher in the western region (average 574, standard deviation 244) compared to the eastern region (average 412, standard deviation 218) (p-value = 0.02838, Supplemental Figure 3A). Shannon diversity followed a similar pattern but was not significant.

### The Mississippi River creates discrete microbiome shifts beyond the expected distance-decay pattern

To evaluate the roles of geographic distance and biogeographic barriers in gut microbiome variation, we subset the data into pairwise comparisons within west, within east, and between regions (east vs west). We found that geographic distance was significantly correlated with both Bray–Curtis (Figure 2A) and Sørensen dissimilarity (Figure 2B). Within the eastern region, Mantel tests revealed significant positive correlations between geographic distance and Bray–Curtis similarity (r = −0.313, *p* = 0.0061, Figure 2A) as well as Sørensen similarity (r = −0.284, *p* = 0.0058, Figure 2B). A similar pattern was observed in the western region, where Bray–Curtis similarity (r = −0.237, *p* = 5.0 × 10⁻⁴, Figure 2A) and Sørensen similarity (r = −0.413, *p* = 1.0 × 10⁻⁴, Figure 2B) were both significantly associated with geographic distance. However, when comparing between regions, Mantel tests revealed a weak but significant correlation between geographic distance and Bray–Curtis similarity (r = −5.536 x 10^8^, *p* =2.41 × 10^-14^, Figure 2A), whereas the relationship between geographic distance and Sørensen similarity was not significant (r = −6.496 × 10^8^, *p* =0.053, Figure 2B). This is partly explained by compositional differences between regions (Supplemental Figure 1), where samples within regions shared about 3.0% of ASVs on average (within west: 3.04%, within east: 3.03%), whereas samples between regions shared only 1.26%. Furthermore, microbial similarity remained significantly lower between regions compared to within regions even when comparing samples within the overlapping distance range (838 km-1492 km, p = < 2.2e-16, Supplemental Figure 2). This demonstrates that geographic distance alone is insufficient to explain the reduced similarity observed across the river. Together, these results suggest that distance–decay of microbiome similarity is not continuous, as evidenced by a discrete shift in microbiome presence and absence patterns between regions compared to within region comparisons.

**Figure 2.**
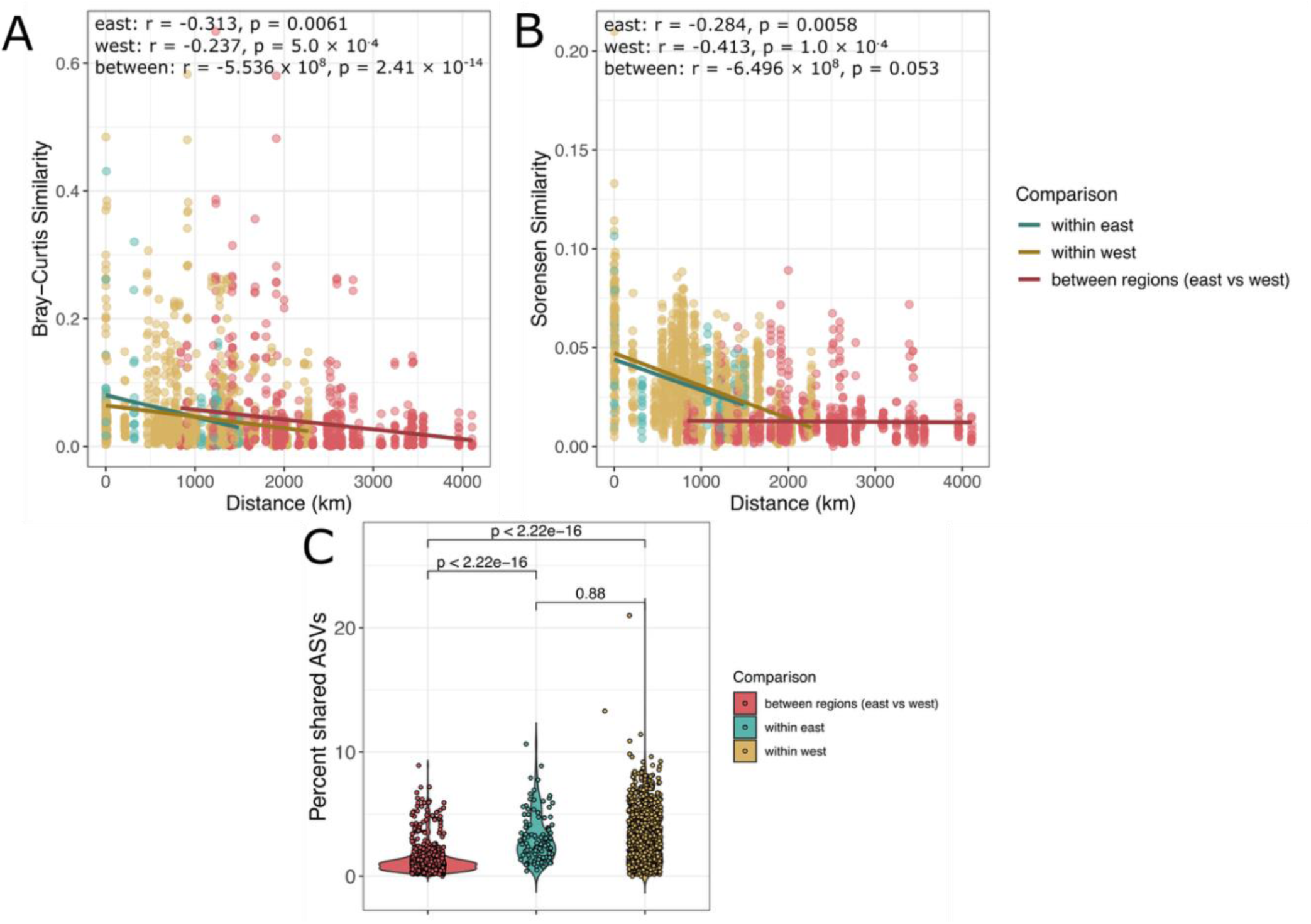
Correlations between microbial measurements vs geographic distance in *Peromyscus maniculatus* fecal samples within and between Mississippi River regions. Comparisons of samples collected in the eastern region are blue, samples collected in the western region are yellow, and comparisons collected from different regions are red. (A) Scatter plot of the correlation between Bray-Curtis Similarity and geographic distance with Mantel test results. (B) Scatter plot of the correlation between Sørensen Similarity and geographic distance with Mantel test results. (C) A violin plot of the percentage of ASVs shared between region (east vs west, red), within east (blue), and within west (yellow) with Wilcoxon test results.

### Bacterial taxa vary in dispersal ability across the Mississippi River barrier

To identify ASVs that were differentially abundant by region, we performed Analysis of compositions of microbiomes with bias correction (ANCOM-BC, Lin & Peddada, 2020). A total of 10 significant ASVs were significantly differentially abundant between east and west (q-value < 0.05, Figure 3A). Of these, four ASVs were assigned to the genus *Lactobacillus*, two were *Helicobacter*, and four were uncultured ASVs (Figure 3A). Five ASVs were enriched in the eastern region and nine ASVs were enriched in the western region (Figure 3A).

**Figure 3.**
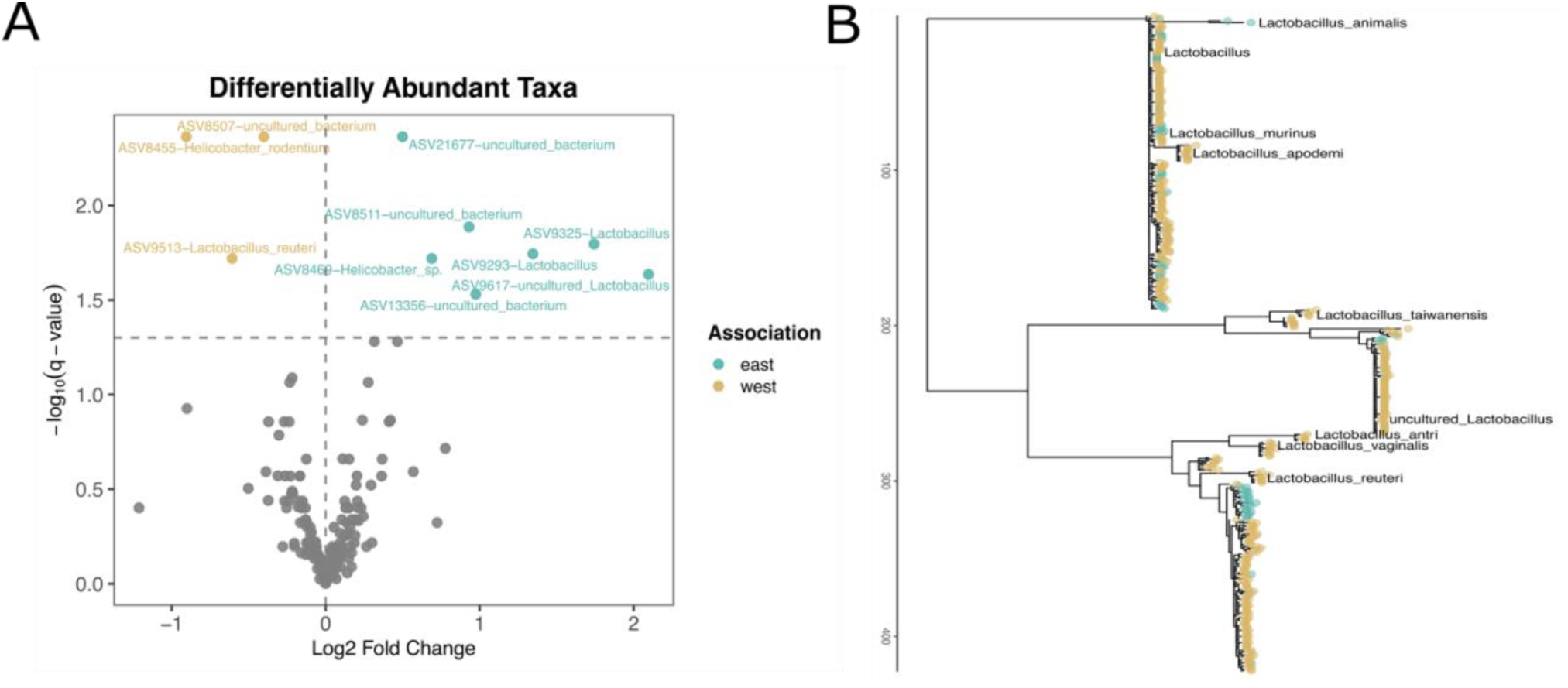
ASVs identified in *Peromyscus maniculatus* gut microbiome associated with either the western or eastern Mississippi region. (A) Significant differentially abundant ASVs (q < 0.5) colored by region (east = blue, west = yellow, gray = insignificant). (B) A phylogenetic tree of genus *Lactobacillus* showing significant clustering of bacterial lineages by region (east = blue, west = yellow, Pagels λ = 0. 972023, p = 7.73E-27, Table 1).

**Table 1.** Phylogenetic clustering of bacterial lineages by region among the top ten most abundant genera identified in *Peromyscus maniculatus*.

| Genus | Lambda<sub>1</sub> | p | # of tips in tree<sup>2</sup> | # differentially abundant ASVs (q val < 0.5) |
| --- | --- | --- | --- | --- |
| *Lactobacillus* | **0.972** | **7.73E-27** | 1655 | 4 |
| *Lachnospiraceae_NK4A136_group* | **0.9739** | **1.59E-287** | 12368 | 1 |
| *Lachnospiraceae_UCG-001* | **0.9685** | **1.70E-19** | 1391 | 1 |
| *Muribaculaceae* | **0.992** | **0.00E+00** | 21262 | 0 |
| *Roseburia* | **0.9733** | **9.83E-29** | 2307 | 0 |
| *Lachnospiraceae_UCG-006* | **0.967** | **2.63E-65** | 1450 | 0 |
| *Colidextribacter* | **0.8053** | **3.62E-03** | 2568 | 0 |
| *Helicobacter* | 0.0001 | 1 | 147 | 2 |
| *Desulfovibrio* | 0.0001 | 1.00E+00 | 1855 | 0 |
| *Ruminococcus* | 0.0001 | 1.00E+00 | 444 | 0 |

<sup>1</sup> Bolded Pagel's $\lambda$ values indicate significance. $\lambda$ values close to 1 suggest a strong phylogenetic signal in which closely related bacterial lineages are clustered by region.
<sup>2</sup> Phylogenetic trees for each genus are shown in Supplemental Figure 4.

To better understand which microbial taxa are more or less sensitive to dispersal across the Mississippi River barrier, we measured the degree of phylogenetic clustering using Pagel’s λ (Pagel, 1999) based on full-length 16S rRNA gene phylogenies (Figure 3B). To obtain sufficient statistical power, we assessed phylogenetic signals within strain-level phylogenies of the 10 most abundant genera (Table 1, Supplemental Figure 4). We found that seven genera were strongly and significantly structured by region (*Muribaculaceae, Lactobacillus, Colidextribacter, Lachnospiraceae_UCG-001, Lachnospiraceae_UCG-006, Roseburia,* and *Lachnospiraceae_NK4A136_group*), whereas three genera showed no evidence of phylogenetic clustering (*Helicobacter, Desulfovibrio, and Ruminococcus*) (Table 1, Supplemental Figure 4). *Lactobacillus* had the highest number of significant differentially abundant taxa (Figure 3A) and was significantly structured by region (Figure 3B).

## Discussion

In this study, we applied ecological concepts traditionally developed for plants and animals (Emerson et al., 2011; Ricklefs & Jenkins, 2011) to assess biogeographic isolation in host-associated microbiomes at a continent-wide scale. Using long-read 16S rRNA sequencing (Buetas et al., 2024; O’Sullivan et al., 2025), we achieved strain-level resolution of the gut microbiota within the *Peromyscus maniculatus* species complex (Figure 3). This represents a technological advancement over traditional short hypervariable 16S rRNA sequencing, particularly for resolving closely related microbial lineages associated with the relatively recent evolutionary divergence of this host system (Boria et al., 2026; Bradley & Lindsey, 2019; Dragoo et al., 2006; Gozashti et al., 2026; Greenbaum et al., 2019). While distance–decay patterns of host-associated microbial communities are well established (Clark et al., 2021; Dillard et al., 2026; Moeller et al., 2017; Suzuki et al., 2019), many non-model systems have focused on comparisons across host species (Chu et al., 2025; Groussin et al., 2017; Kohl et al., 2022; Li et al., 2025; Tuoliu et al., 2025) rather than within a single species. Here, we demonstrate a striking regional geographic signal associated with microbial variation (Figure 1A, 1B) and distance decay across the continent within the *P. maniculatus* species complex (Figure 2A, 2B). This framework allowed us to explicitly test whether the Mississippi River constrains microbial assembly and transmission and whether the observed dissimilarity exceeds expectations based on geographic distance alone.

We identified three major patterns in the biogeographic distribution of the *P. maniculatus* gut microbiome. First, we observed a continuous distance-decay relationship within both the east and west regions across Bray-Curtis and Sørensen distances (Figure 2A, 2B). This indicates that within a single side of the river, microbial dispersal is continuous but limited by distance, with closer hosts sharing more similar community composition and relative abundances. Second, when comparing microbial composition between the east and west regions, we found little to no evidence of a distance-decay relationship using the Sørensen index (Figure 2B). This suggests that the Mississippi River acts as a strong barrier to microbial colonization, creating a discontinuity in the regional species pool, likely due to the river limiting host movement. In contrast, Bray-Curtis distance showed a continuous distance-decay relationship between east and west sites (Figure 2A), whereas Sørensen distances exhibited a constant plateau regardless of geographic distance (Figure 2B). This can be explained by species turnover on each side of the river, while their relative abundances continue to shift along a continental-scale environmental gradient. Third, our strain-level analysis identified microbial taxa that vary in their sensitivity to this biogeographical barrier. For some genera, taxa differ in both abundance and strain identity between east and west, further supporting the idea that the river acts as a barrier to strain transmission and results in turnover of microbial taxa.

Geographic distance often covaries with other biologically relevant factors, particularly host genetic distance (Andreu-Sánchez et al., 2025) and environmental dissimilarity (Groussin et al., 2020). If microbiomes are partially structured by host-associated traits, then increased host divergence could drive divergence in microbial communities (Couch et al., 2020). Host populations distributed across environmental landscapes may experience both genetic divergence and differing ecological conditions, reinforcing microbiome differentiation (Couch & Epps, 2022). Our results suggest that host isolation across the Mississippi River amplify microbiome differences beyond what would be expected from geography and ecology alone (Tung et al., 2015). However, in the *P. maniculatus* species complex, geographic separation appears to be a stronger driver of gut strain diversity than maternal transmission (Dillard et al., 2026), indicating that spatial processes often outweigh microbial inheritance within host species. Notably, the California population had the most divergent microbiome in our study populations (Figure 1A, Supplemental Figure 1), raising the possibility that the Sierra Navada acts as an additional geographic barrier limiting microbial dispersal. Interestingly, Boria et al. (2026) showed no evidence of reduced host gene flow within western populations of *P. m. sonoriensis*. This decoupling suggests that the gut microbiome may serve as a sensitive bio-indicator of ecological connectivity, potentially capturing finer-scale landscape or habitat features that are not yet apparent in host nuclear genetic markers. Such findings imply that while taxa may be broadly shared across space, their relative dominance is shaped by additional mechanisms such as dispersal barriers, historical range shifts, or localized environmental variation that remain uncaptured by models of distance decay alone.

We leveraged the resolution of full-length 16S rRNA gene phylogenies to assess the influence of biogeographic barriers on bacterial dispersal. The regional specificity of microbial taxa differed among bacterial genera, indicating that geographic barriers do not uniformly restrict microbial exchange but instead differentially affect taxa depending on their ecological and evolutionary traits. Genera exhibiting strong regional structure, including *Lactobacillus*, *Lachnospiraceae_UCG-001*, and *Lachnospiraceae_NK4A136_group* (Table 1), represent taxa with limited dispersal and high host fidelity (Frese et al., 2011; Walter, 2008). Additionally, these bacteria are host-associated colonizers that persist within the gut (Meehan & Beiko, 2014; Vacca et al., 2020). Such traits would reduce opportunities for transmission between populations, thereby promoting regional lineage diversification. At the opposite end of the spectrum, genera such as *Desulfovibrio* and *Ruminococcus* showed neither phylogenetic structure nor differential abundance (Table 1), suggesting minimal barriers to dispersal and high connectivity between regions. Both taxa play essential roles in the gut microbiome compared to other taxa. Specifically, *Desulfovibrio* is the most predominant genera of sulfate-reducing bacteria (Rey et al., 2013) and *Ruminococcus* degrade and convert complex polysaccharides into a variety of nutrients for the hosts (La Reau & Suen, 2018). However, the homogenous distribution of these taxa could also reflect generalists, where these taxa are broadly distributed in the environment and readily colonize hosts regardless of region (Chen et al., 2021).

While we were unable to resolve whether ecological similarity plays a part in the river effect seen here, our results highlight that processes beyond geographic distance shape microbiome composition at large scales. Future research could utilize arboreal habitats on opposite sides of the continent as a powerful natural experiment to control for environmental similarity with increased sampling efforts (Bedford & Hoekstra, 2015; Boria et al., 2026; Hager & Hoekstra, 2021). To explicitly test the relative contributions of host dispersal versus microbial traits in transmission, future work should integrate host genomic sequencing with shotgun metagenomics. Comparing multiple host species with differing dispersal capacities could further clarify the generality of these patterns at deeper evolutionary timescales. Finally, to determine whether bacterial strain-level divergence reflects functional differentiation, future studies could incorporate metagenome-assembled genomes to compare the functional gene content of shared microbial species between regions, similar to approaches in human cohorts (Tett et al., 2019).

Overall, we showed that the *P. maniculatus* species complex exhibits: (i) distance-decay relationships within both the eastern and western regions using Bray-Curtis and Sørensen distance, (ii) species turnover between regions that exceeds expectations based on geographic distance alone, and (iii) variable sensitivity among bacterial genera to this biogeographic barrier, finding enabled by our strain-level phylogenetic analysis of full-length 16S rRNA sequences. Broadly, our findings suggest that microbiome assembly is governed by the interplay of dispersal constraints and landscape features, highlighting the need to look beyond spatial models to predict microbial biogeography.

## Acknowledgments

The National Ecological Observatory Network (NEON) is a program sponsored by the National Science Foundation and operated under a cooperative agreement by Battelle. Laura Steger and the NEON Biorepository at Arizona State University (ASU) genotyped the samples, assisted with sample selection, and provided the samples and associated data collected through the NEON Program. We thank Lina Ali for assistance with DNA extraction of rodent fecal pellets and the Research Computing at ASU for providing computing and data storage resources. Finally, we thank members of the ASU Biodesign Center for Health through Microbiomes for helpful comments and support during the development of this project. T.A.S. was supported by the National Institutes of Health (R35 GM160076).

## Availability of data and materials

The raw reads have been deposited in SRA under. All R scripts used in this project are available through GitHub at https://github.com/DaniBlumstein/pema_NEON.

## Authors’ contributions

Conceptualization: D.M.B., T.A.S; Methodology: D.M.B., G.S.,T.A.S; Formal analysis: D.M.B.; Resources: T.A.S.; Data curation: A.P., D.M.B., T.A.S.; Writing - original draft: D.M.B.; Writing - review & editing: D.M.B., G.S., A.P., T.A.S.; Visualization: D.M.B.; Supervision: T.A.S.; Project administration: T.A.S.; Funding acquisition: T.A.S.

## Supplemental

**Supplemental Table 1.** Selected samples from the National Ecological Observatory Network (NEON) as part of the small mammal box trapping protocol of *Peromyscus maniculatus*.

**Supplemental Table 2.** Sampling totals of *Peromyscus maniculatus* by state and sex.

| State | Females | Males | Total | 635 |
| --- | --- | --- | --- | --- |
| California | 1 | 2 | 4 | 636 |
| Colorado | 5 | 6 | 11 | 637 |
|  |  |  |  | 638 |
| Kansas | 2 | 3 | 5 | 639 |
|  |  |  |  | 640 |
| New Hampshire | 2 | 1 | 3 | 641 |
|  |  |  |  | 642 |
| North Dakota | 3 | 3 | 6 | 643 |
|  |  |  |  | 644 |
| Oklahoma | 3 | 3 | 6 | 645 |
|  |  |  |  | 646 |
| Tennessee | 2 | 2 | 4 | 647 |
|  |  |  |  | 648 |
| Utah | 2 | 4 | 6 | 649 |
|  |  |  |  | 650 |
| Virginia | 1 | 3 | 4 | 651 |
|  |  |  |  | 652 |
| Washington | 3 | 3 | 6 | 653 |
|  |  |  |  | 654 |
| Wisconsin | 2 | 3 | 5 | 655 |
|  |  |  |  | 656 |
| Wyoming | 3 | 3 | 6 | 657 |
|  |  |  |  | 658 |

**Supplemental Table 3.** PERMANOVA results of regional effects on the microbiome accounting for latitude and longitude at three microbial taxonomic levels and two distance metrics.

| Level | Metric | R<sup>2</sup> | F | P value |
| --- | --- | --- | --- | --- |
| ASV | Bray-Curtis | 0.0744 | 1.6618 | 0.01 |
| ASV | Sørensen | 0.0717 | 1.5957 | 0.01 |
| Species | Bray-Curtis | 0.0987 | 2.2634 | 0.01 |
| Species | Sørensen | 0.1093 | 2.5357 | 0.01 |
| Genus | Bray-Curtis | 0.1615 | 3.9799 | 0.01 |
| Genus | Sørensen | 0.1225 | 2.8853 | 0.01 |

---

**Supplemental Figure 1.**
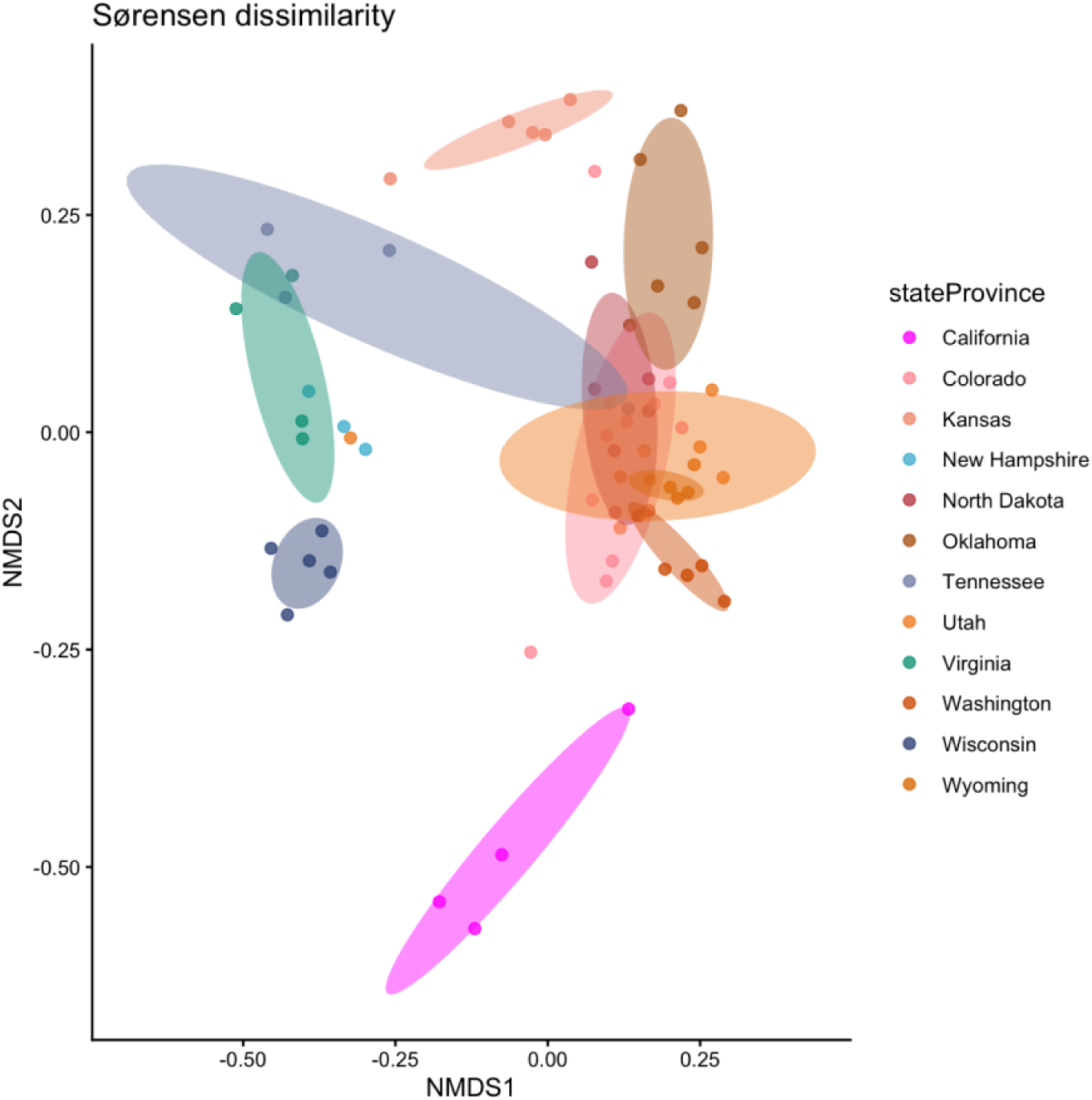
NMDS plot showing the effect of collection location on Sørensen dissimilarity. Colors correspond to the state the samples were collected in.

**Supplemental Figure 2.**
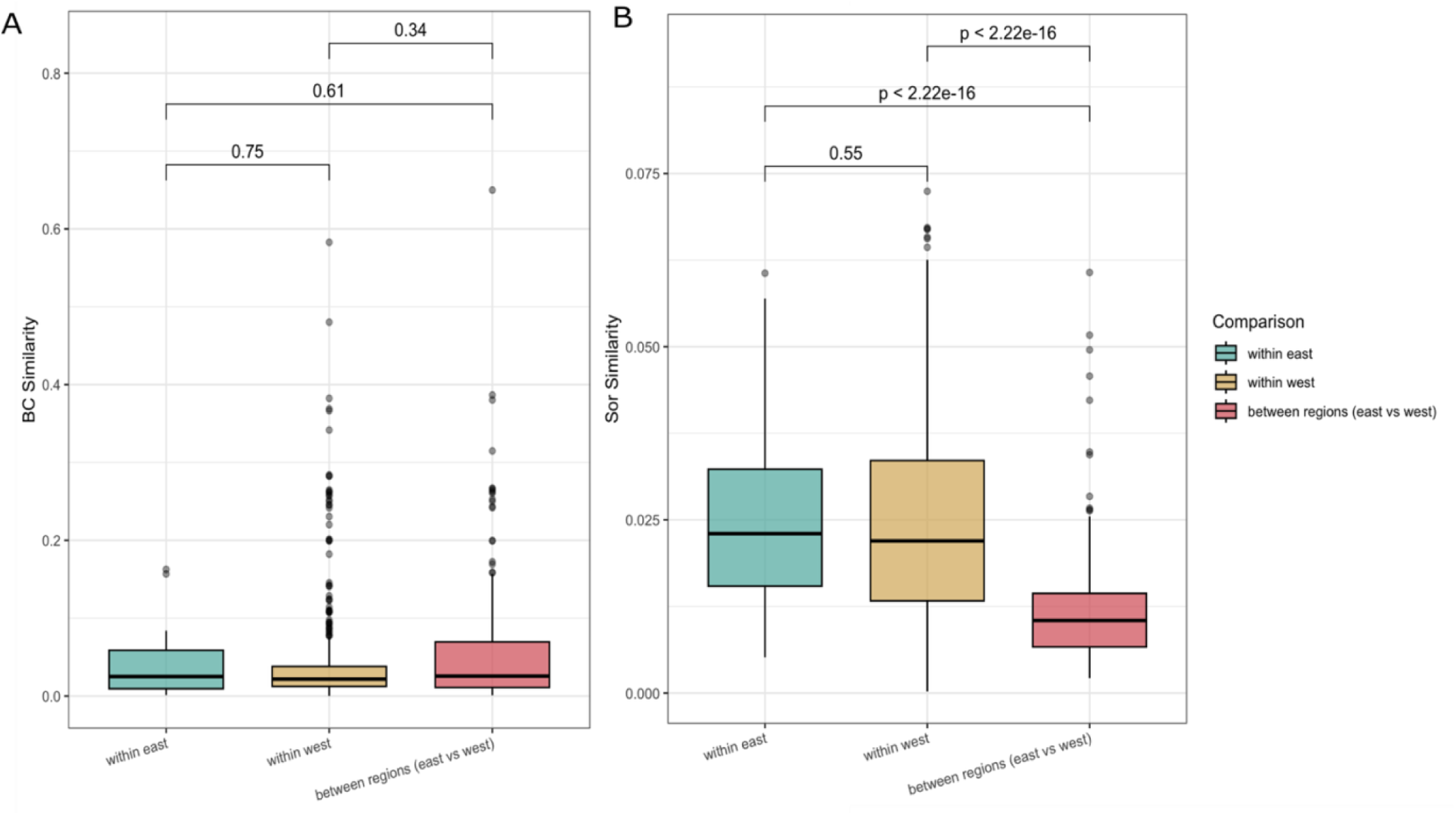
Comparison of beta diversity within versus between regions only across the overlapping distance range (838 km-1492 km). Samples collected in the eastern region are blue, samples collected in the western region are yellow, and comparisons collected from different regions are red. (A) Box plot of Bray-Curtis Similarity and geographic region with Wilcoxon test p value results. (B) Box plot of Sørensen Similarity and geographic region with Wilcoxon test p value results.

**Supplemental Figure 3.**
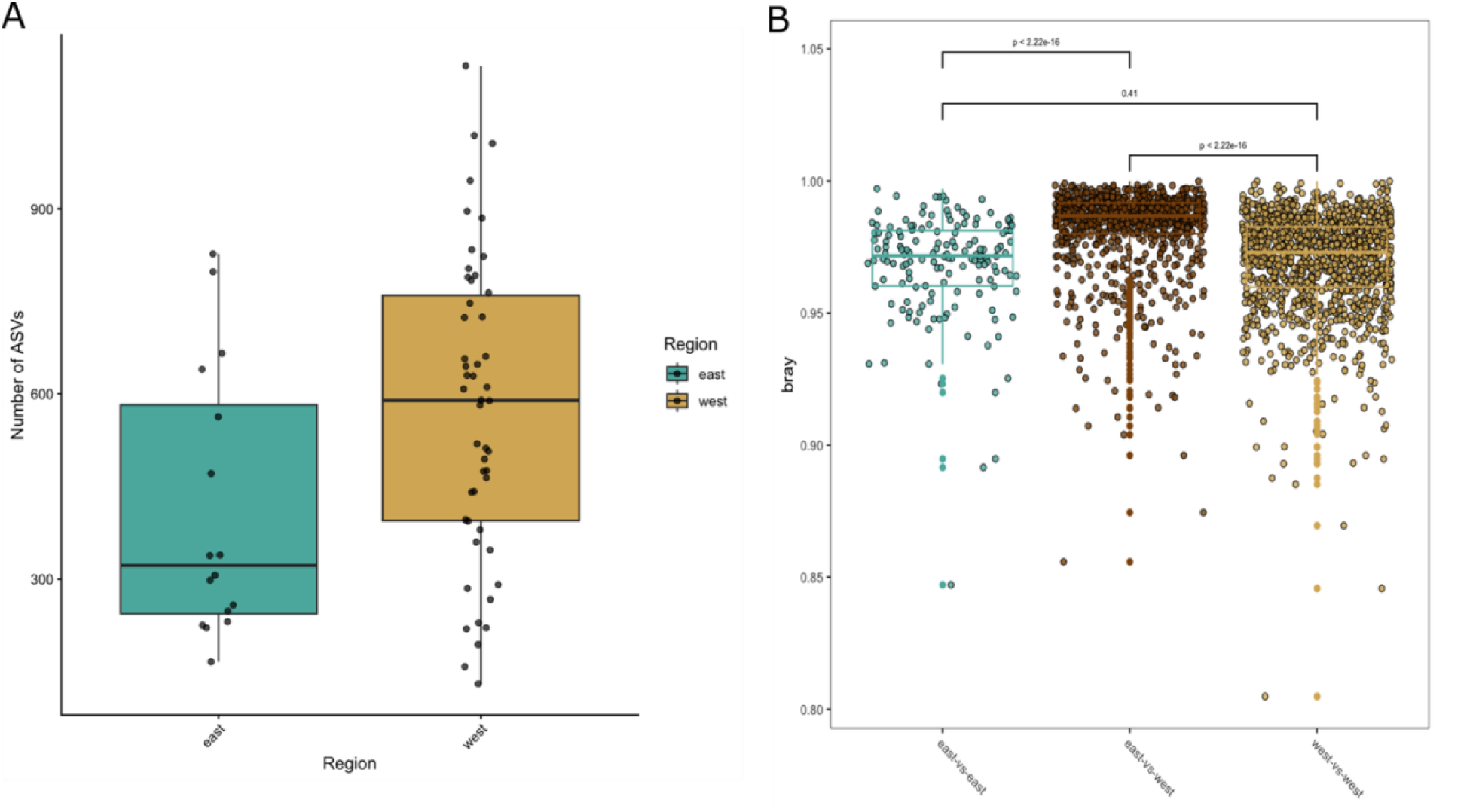
Differences in ASV composition of *Peromyscus maniculatus* gut microbiome between Mississippi river regions. Samples collected in the eastern region are blue, samples collected in the western region are yellow, and comparisons for east vs west are brown. (A) Box plot the number of ASV by geographic region. (B) Box plot of pairwise comparison of Bray-Curtis distance among the geographic regions with Wilcoxon test p value results.

**Supplemental Figure 4.**
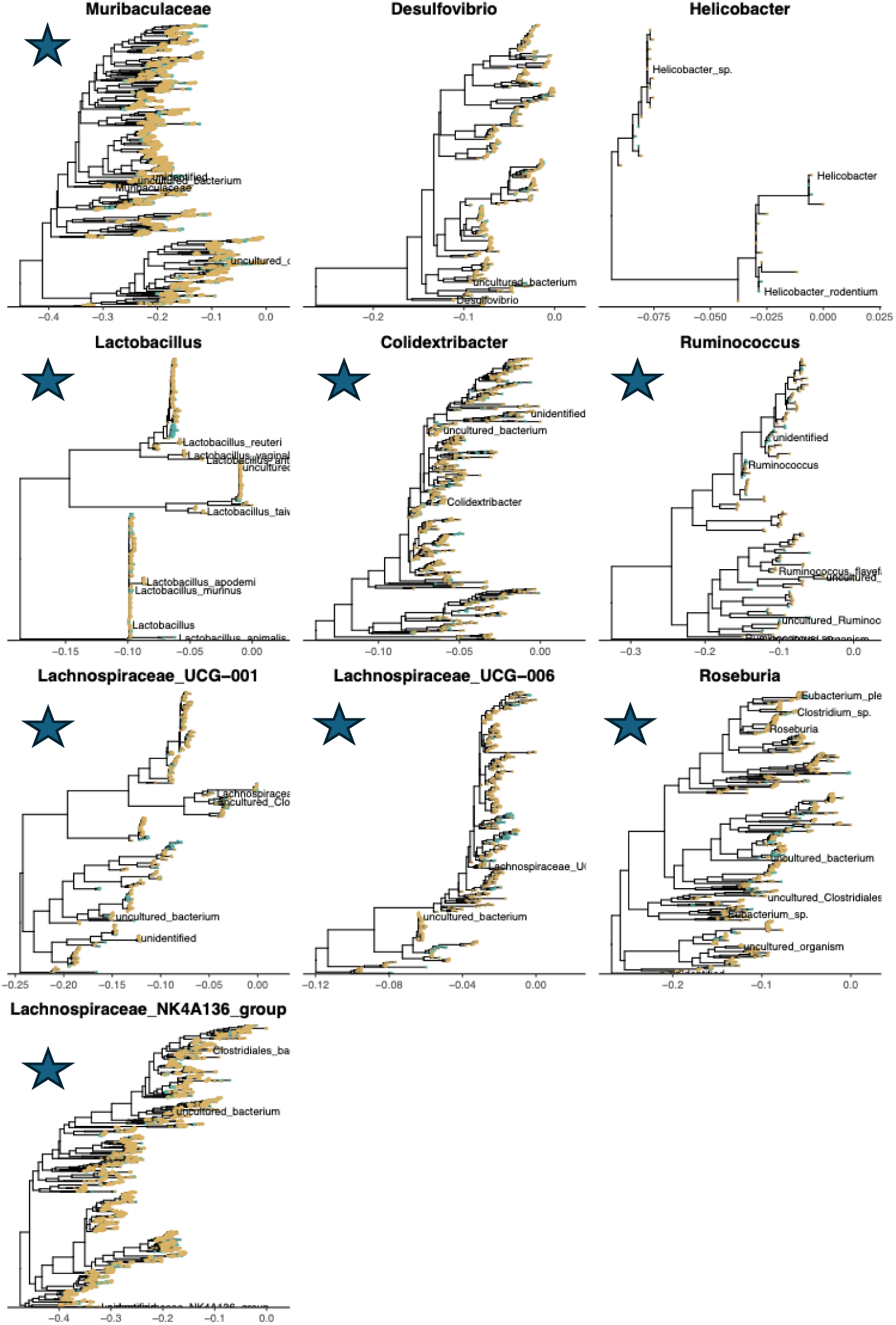
Top ten genera in the *Peromyscus maniculatus* gut microbiome dataset. ASVs found on the western side of the Mississippi River are colored yellow while ASVs found on the eastern side of the Mississippi River are colored blue. Stars are genera that are significant for lambda.

## Notes

https://github.com/DaniBlumstein/pema_NEON

### Competing Interest Statement

The authors have declared no competing interest.

